# Structural analysis and ensemble docking revealed the binding modes of selected progesterone receptor modulators

**DOI:** 10.1101/2022.10.06.511110

**Authors:** F. Saritha, N. Aiswarya, R. Aswath Kumar, K.V. Dileep

## Abstract

Uterine fibroids (UF) are benign smooth muscle neoplasm of uterus that have a significant impact on a woman’s quality of life as they perturb hormonal homeostasis resulting in heavy menstrual bleeding, impaired fertility, pregnancy complications and loss. UF can be surgically removed through invasive procedures, but their recurrence rate is often high. Progesterone receptor (PR) has an imperative role in UF management. Mifepristone, ulipristal acetate (UPA) and asoprisnil (ASO) are some promising selective progesterone receptor modulators (SPRMs), acts on PR, but due to their side effects in long term use, they were withdrawn from the market. Hence, there is a dire need for novel, highly efficient with least side effects, therapeutics for the treatment of UF. To contribute towards the drug discovery for UF, we made an extensive structural comparison of reported PR crystal structures, also elucidated the binding modes of four existing SPRMs against human PR through ensemble docking approach. Our studies revealed the presence of 5 highly repeating water molecules that has an important role in ligand binding and structural stability. Our ensemble docking and MD simulation revealed that studied ligands have preferential selectivity towards the specific conformation of PR. It is anticipated that our study will be a useful resource to all the drug discovery scientists who are engaged in the identification of lead molecules against UF.

## Introduction

Uterine fibroids (UFs), also known as uterine leiomyomas, are benign smooth muscle neoplasms of the uterus that affect women of reproductive age [1]. The UFs can often cause a wide range of severe and chronic symptoms like heavy menstrual bleeding, which can lead to anaemia, fatigue and painful periods [2]. Sometimes UFs may be associated with reproductive problems, including impaired fertility, pregnancy complications and loss, and adverse obstetric outcomes [3]. Women manifested with UF in near menopausal age of 46-50 years is around 62.8%, when compared to 21.3% in 30-35 years. The risk factors of UFs include age, race, endogenous and exogenous hormonal factors, obesity, uterine infection, and lifestyle (diet, caffeine and alcohol consumption, physical activity, and stress) [4]. Although, UF can be surgically removed through invasive procedures such as myomectomy, laser ablation, or embolization to improve the fertility; the recurrence rate is often high. Therefore, an unmet medical need is there for non-invasive treatment of UF.

Since ovarian steroid hormones such as estrogen and progesterone play an important role in the growth of UF [5], manipulation of these hormones mediated signalling has become a key treatment strategy. Non-invasive treatment approaches include Gonadotropin releasing hormone (GnRH) receptor agonists and antiprogestins. But GnRH agonist can cause severe menopausal symptoms, and cannot be used for long-term treatment [6]. Mifepristone (Figure 1A), also known as RU486, acts through the inhibition of progesterone receptors (PRs), which seems to have a crucial role in the growth of UFs [7]. Studies revealed that mifepristone reduced UF-associated bleeding and improved fibroid-specific quality of life, without reducing fibroid volume [8].

**Figure 1:**
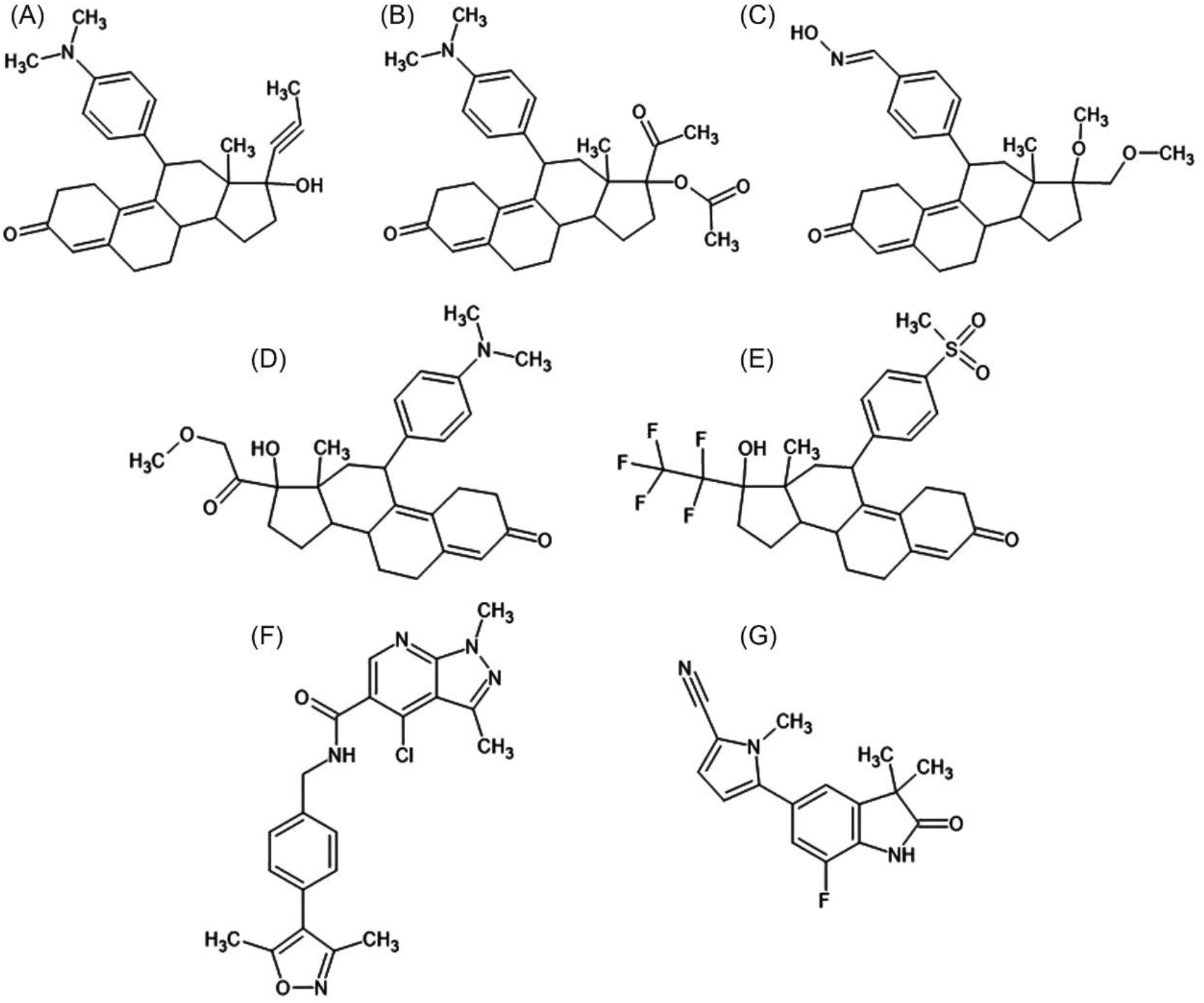
Chemical structures of selected PR modulators: (A) mifepristone, (B) ulipristal acetate, (C) asoprisnil, (D) Telapristone, (E) Vilaprisan, (F) GSK-1564023A (G) WAY-255348.

The specific progesterone receptor modulators (SPRMs) have been emerged as an alternate and promising therapeutic agent for UF, endometriosis and breast cancer [9]. Both biochemical and clinical evidence suggests that SPRMs may reduce fibroid growth and ameliorate symptoms [8]. Based on the chemical compositions, the SPRMs and antagonists were broadly classified into two categories like steroidal and non-steroidal molecules. The action of SPRMs is tissue-specific, leading to both agonistic and antagonistic effects [10]. SPRMs such as ulipristal acetate (UPA) (Figure 1B) and asoprisnil (ASO) (Figure 1C) decrease extracellular matrix formation and have been efficiently used for the management of UF [11]. Another SPRM, Telapristone (also known as Proellex)(Figure 1D) has been evaluated for the symptoms associated with endometriosis and UF [12]. Vilaprisan (BAY 1002670) (Figure 1E) is a new orally available SPRM, with anti-proliferative activity against UFs [10]. However, due to the safety concerns, many FDA approved SPRMs (such as UPA and ASO) were withdrawn from the market. ASO causes unfavorable endometrial changes in UF patients and UPA was deemed to have liver toxicity [11, 13]. Hence there is a dire need for new drug with high efficacy and little side effects.

The computer-aided drug discovery (CADD) became a powerful, and promising approach for faster, cheaper, and more effective ways of drug design. The rapid growth of computational tools for drug discovery has exhibited a significant and outstanding impact on anticancer drug design, and has also provided fruitful insights into the area of cancer therapy [14,15]. Hence, CADD is certainly important for the identification of novel lead molecules, which will be useful to stop the disease progression [16, 17].

In the present study we explained the binding modes of selected SPRMs/antagonist having steroidal (Telapristone and Vilaprisan) and non-steroidal scaffolds (Figure 1F and G) (GSK-1564023A and WAY-255348) against human PR through ensemble docking approach. It is anticipated that the current study will be useful to the computational scientists to discover novel SPRMs/ antagonists for progesterone receptors.

## Materials and Methods

### Selection and Preparation of Progesterone Receptors from PDB

To study the heterogeneity in the ligand binding patterns in various crystal structures of human PR, we collected all reported structures from PDB (www.rcsb.org) and compared each other, especially their binding sites were critically analyzed. Since our focus was on the antagonists and SPRMs bound human PRs, we excluded agonist bound PRs. We then grouped the collected structures into four categories based on the steroidal scaffold present in the ligands. The four groups are as follows, antagonists (Group-1) and SPRMs (Group-2) that contains steroidal scaffold, antagonists (Group-3) and SPRMs (Group-4) that consists of non-steroidal scaffolds. The high-resolution structures from each group were selected as candidate structures for further studies. These selected structures were prepared using protein preparation wizard of Maestro, Schrödinger suite. During preparation, the protonation, bond order, disulphide bridges, orientations of the amino acids were corrected. Polar hydrogen atoms were added to the protein and these hydrogen atoms were optimized. A short energy minimization with a cut-off of 0.30 Å was also performed. All the receptor structures were prepared in the presence and absences of invariant water molecules [18, 19].

### Identification of highly repeating water molecules

Using a web based interactive computing server 3dSS (http://cluster.physics.iisc.ernet.in/3dss/) we identified all the invariant water molecules in the crystal structures of human PR. We selected 18 crystal structures of human PR from PDB and was superposed to a fixed molecule, PDB ID: 1SQN using ProFit algorithm and identified highly repeating water molecules within 2.5Å of binding site [20].

### Selection and preparation of Ligands

We identified several steroidal and non-steroidal antagonists and SPRMs from the literature and downloaded their structures from different database like PubChem (https://pubchem.ncbi.nlm.nih.gov/), clinical trial.gov (https://www.clinicaltrials.gov/) and Therapeutic Target Database (http://db.idrblab.net/ttd/). Ligands that are not available in these databases were drawn using 2D sketcher tool of Maestro, Schrödinger suite [21]. Selected ligands were then prepared using Ligprep module of the Schrödinger suite. During the ligand preparations, various conformations of the ligand at pH 7±2 was generated and used for our studies.

### Molecular Docking

To intuit the binding mode of different known antagonists and SPRMs towards the human PRs, we performed ensemble docking using Glide (standard precision docking) [22]. The contributions of highly repeating water molecules in the ligand binding were also assessed through docking studies by including them in our calculations. The binding energies of the ligands were also calculated using prime MM-GBSA module of Schrödinger suite [23].

### Analysis of interactions

In order to investigate binding mode of selected ligands at the ligand binding domain (LBD) of the PR, we tabulated all types of ligand-interactions with the protein using Matrix plot, Asteroid plot available in Protein Contacts Atlas (https://www.mrc-lmb.cam.ac.uk/rajini/index.html), Pymol [24], and Maestro, Schrödinger suite.

### Molecular Dynamics Simulation

Molecular dynamics (MD) simulations using GPU Desmond [25] was carried out to understand the conformational flexibility of PR LBD upon ligand binding. We used the docked poses for the MD simulation. The system for MD simulation was prepared by soaking them into an orthorhombic box having TIP3P water molecules. The whole system was neutralized with appropriate number of counter ions (We added Na+ or Cl-atoms). The highly repeating 5 water molecules were also retained in the MD simulations. The duration of the MD simulation was 100 ns and the intermediate structures and energies were recorded at every 100 and 1.2 ps respectively.

## Results and Discussions

### Hypothetical clustering of PR structures revealed 4 groups

In structure-based drug discovery, the structural heterogeneity of residues located in the ligand binding site plays an important role in the identification of novel ligands and often helps in the prediction of accurate binding modes. The structural heterogeneities present in the available PRs has been assessed by comparing them. We retrieved all the 18 crystal structures of human PR from PDB (Table 1). Among 18 crystal structures, only 12 structures were of our interest as they are complexes with small molecules such as antagonist and SPRMs. We excluded 6 agonist bound crystal structures. The selected 12 structures were further clustered in to 4 groups depending on the scaffold present in each ligand. The group-1 consists of steroidal antagonists. On the other hand, the group 2 to 4 consists of steroidal SPRMs, non-steroidal antagonists and non-steroidal SPRMs respectively (Table 1).

**Table 1:**
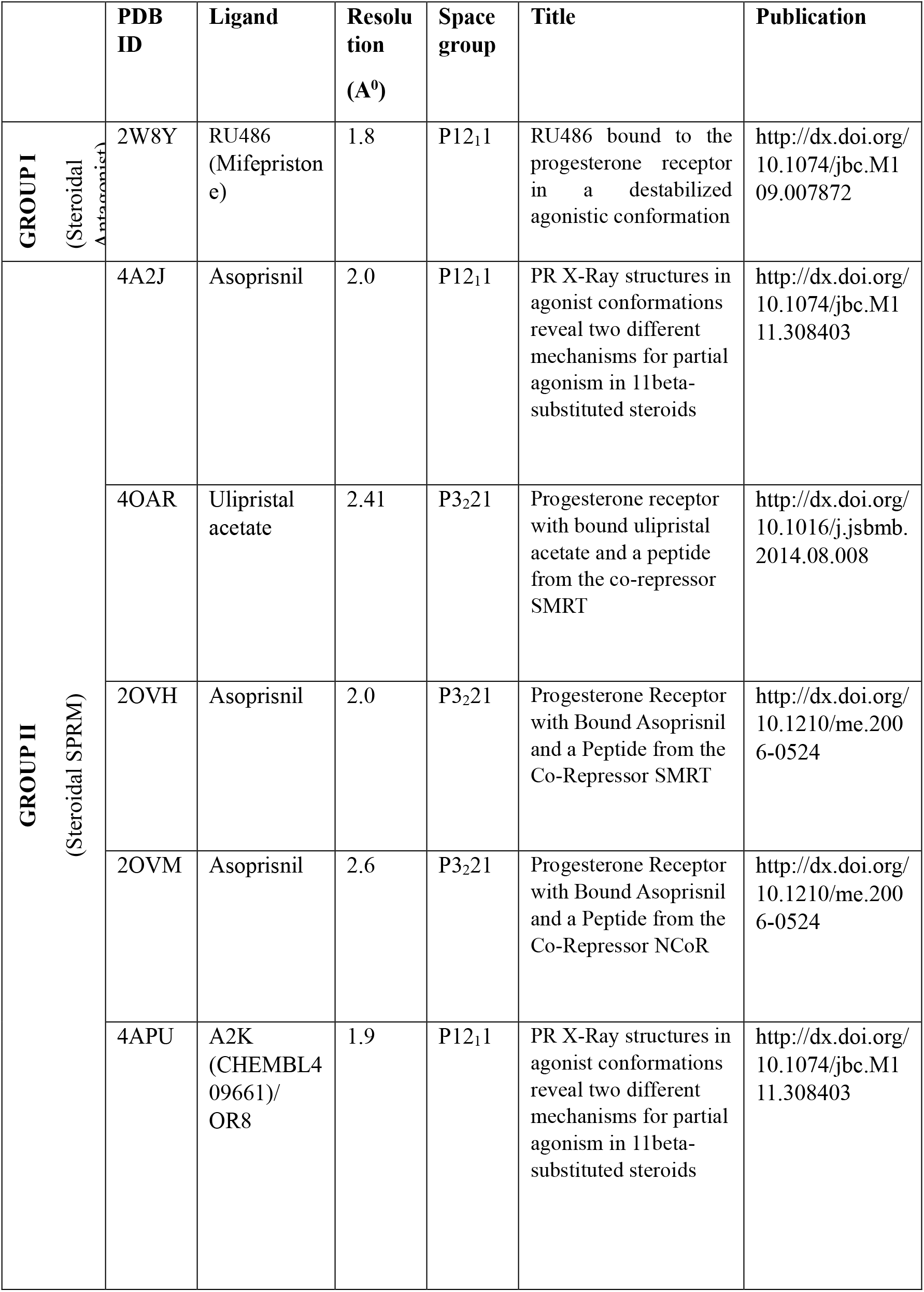
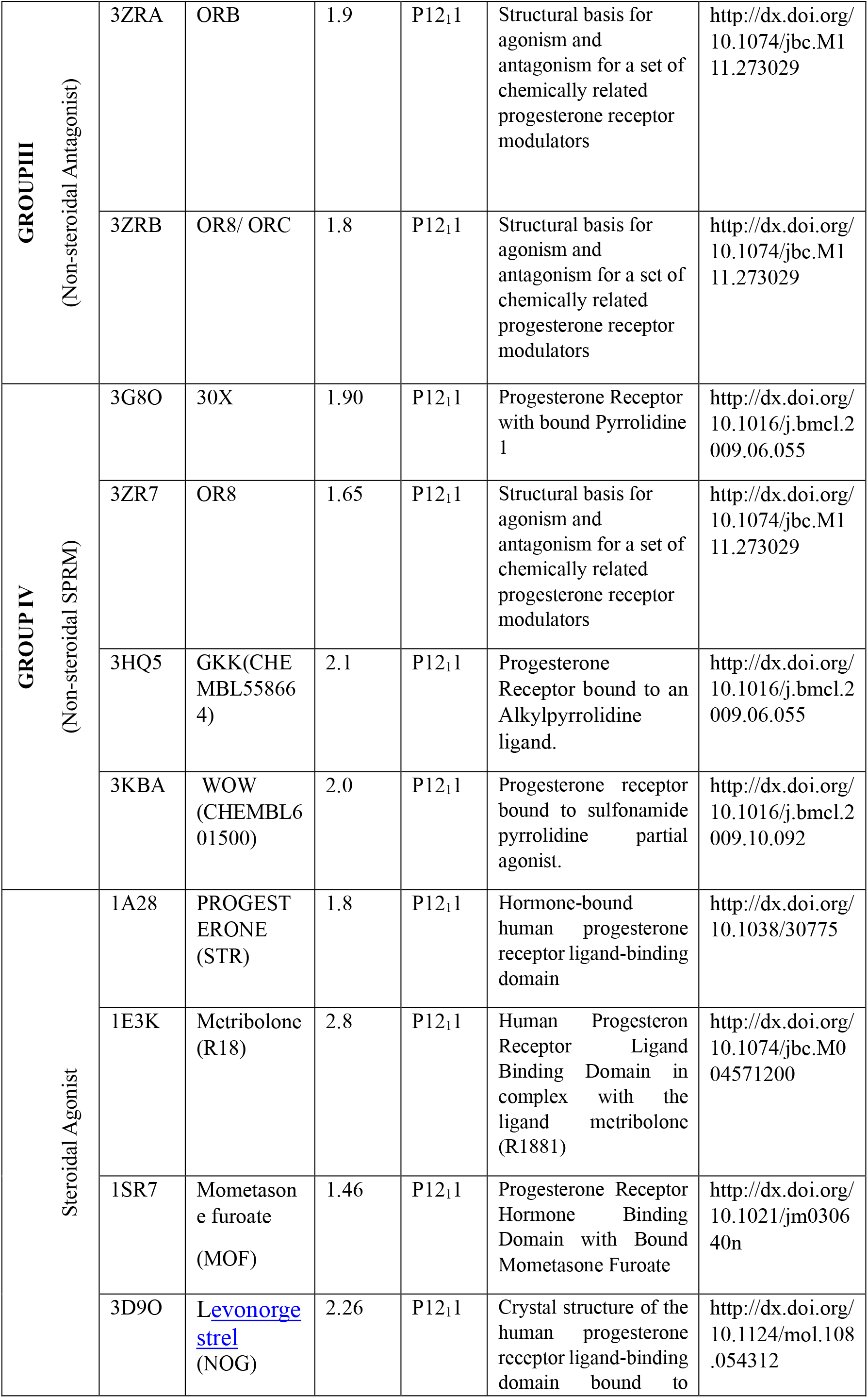
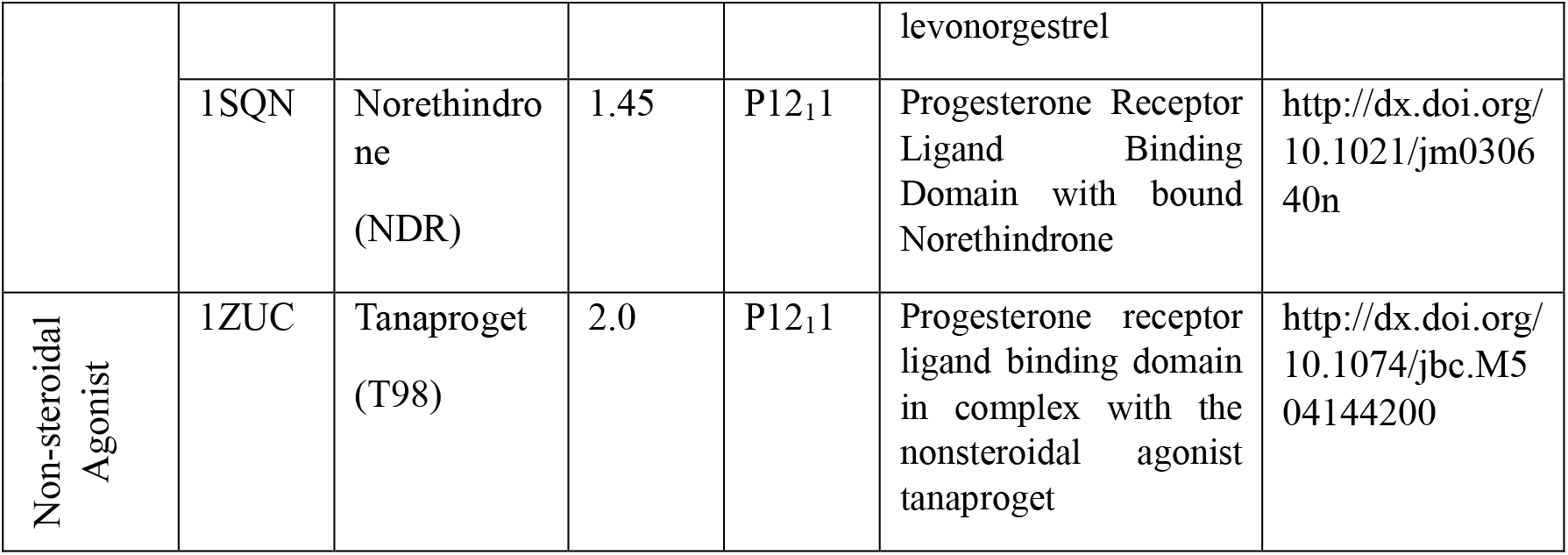
Details of all 18 human progesterone receptors structures retrieved from PDB database. Out of these, 12 antagonists and SPRM bound structures were extensively used for the present study and their classification based on the nature of ligands bounds were as follows.

### PR consists of several highly repeating water molecules

Our analysis to find out invariant water molecules revealed several highly repeating water molecules. The ligand binding domain of PR in complex with norethindrone (PDB ID 1SQN) was used as a reference structure as it has the best resolution and all other proteins were superimposed to the reference molecule. We used 2.5 Å as a probe radius to identify water molecules. Our analysis identified 5 highly repeating water molecules those were scattered around the protein (named Wat-1 to Wat-5 respectively) (Figure 2), in which Wat-1 (Figure 3A) to 3 (Figure 3B) were located near to the loop region of PR after helix-1 (H1). The Wat-4 has interactions with Q725, R766 and with the ligands (both steroidal and non-steroidal ligands) (Figure 3B). Finally, Wat-5 is located in the proximity of the loop between H8 and H9 helix of the PR (Figure 3C). Wat-1 and 2 repeats in 88.23% human PR structures whereas Wat-5 in 82.35% structures and Wat-3 in 76.47% (Figure 3D). Studies have reported that there is a conformational change for the loop located between helix-1 and 3 in the unliganded and liganded LBDs structures [26], and we assume that such kind of conformational changes are partly mediated through the highly repeating water molecules located at the vicinity of the loop. These repeating water molecules may also help in the entry of ligands into LBD of proteins [27]. Wat-4 plays an important role in ligand binding, as it is having a direct hydrogen bonding with the PR, also this water molecule is conserved in all the 18 crystal structures of human PR [27-29]. The position of Wat-5 is in such a way that it is trapped inside the protein, strongly contributes to the structural stability of PR.

**Figure 2:**
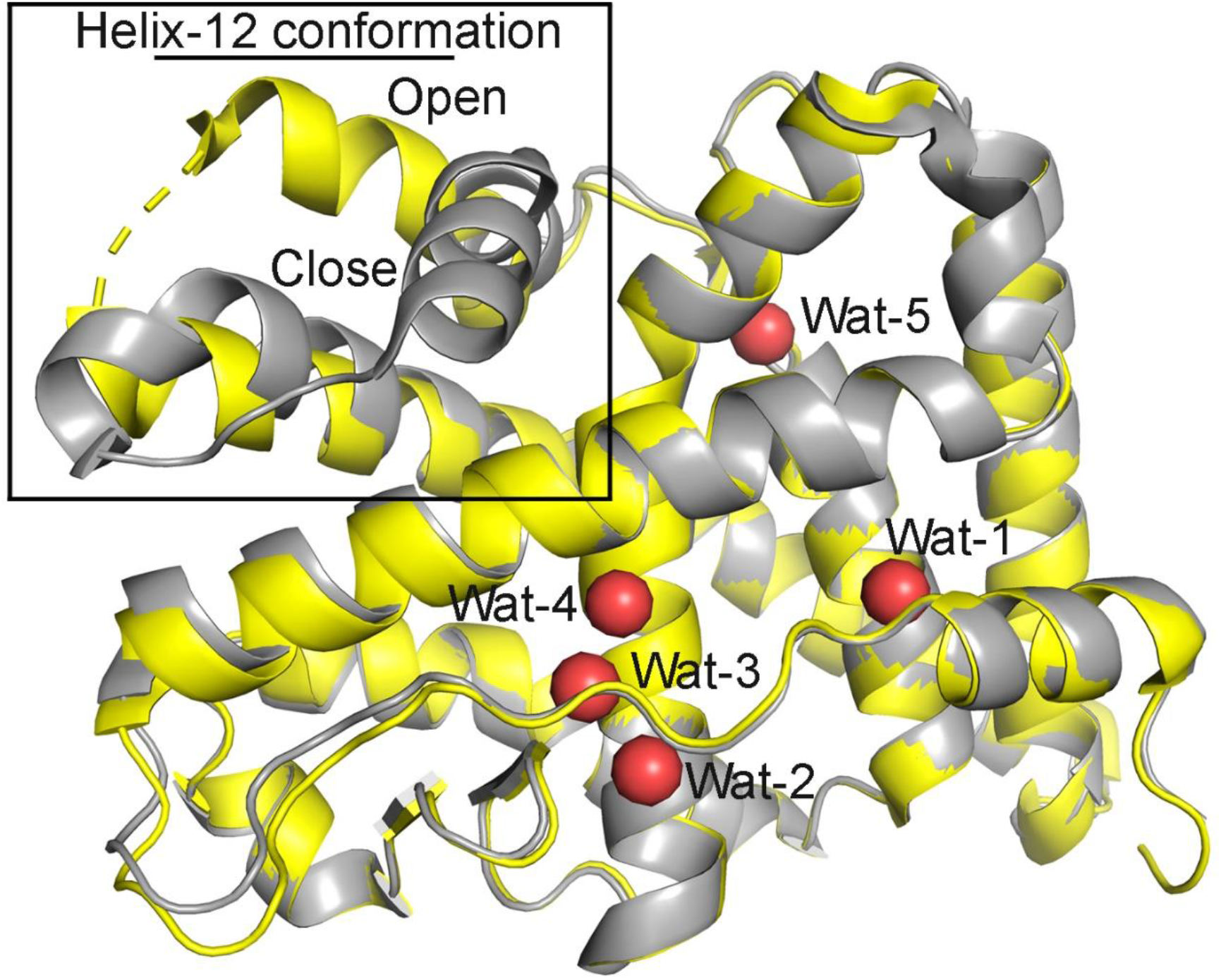
Position of five highly repeating water molecules found in the 18 crystal structures of human PR. Two different conformations (open and close) available for the LBD is also mentioned.

**Figure 3:**
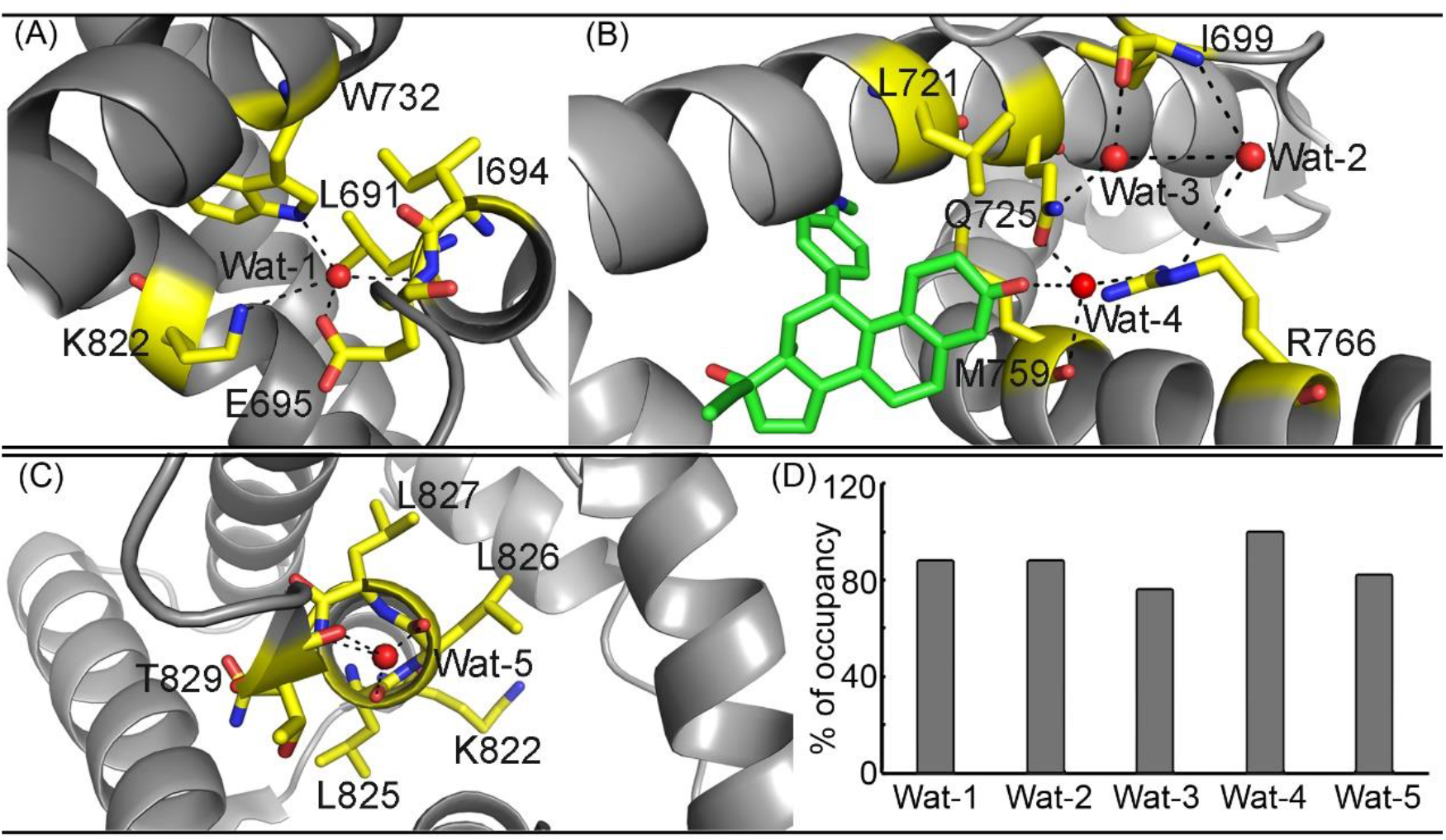
Interaction of five highly repeating water molecules with PR: (A) the hydrogen bonding pattern of Wat-1 (A), Wat 2-4 (B) and Wat-5 with the residues in PR. Water molecules were shown as red spheres and the protein atoms were shown as thick stick and cartoon model. Hydrogen bonding were represented as dashed lines. (D) The percentage of occupancy of highly repeating water molecules in 18 crystal structures of human PR.

### Plasticity of human PR structures

When we closely analyzed the available crystal structures of human PR in the PDB, it was noticed that the B-factors for loops located between H1 and H3, H9 and H10, and H11 and H12 were higher in SPRM bound structures when compared to antagonist and agonist bound structures [30-33]. These loops were already referred as highly flexible regions in the previous studies [33]. i.e., the plasticity of the protein is largely determined by the ligands binds to the LBD. When a steroidal SPRM binds to the PR, a conformational flexibility was observed to the loop that connects helix 12 (H12) [near to M909]. This conformational flexibility may be due to the bulky C11 substitution of SPRM, which protrudes from the binding pocket causes clash with M909, results in the shift at the loop region [34]. This movement provide space for the longer corepressor helix and also displaces E911, which is critical for co-activator binding [30,35]. Conversely, when a non-steroidal SPRM binds to the PR, plasticity was observed in the regions from 703-710 and 859-861.

Another observation in the case of antagonistic human PR structures, was that the co-repressors fabricate multiple polar interactions with residues R740, K734 and E752 [30]. And these interactions may account for the strong recruitment of co-repressor by antagonistic ligand [30].

### Ensemble docking of selected PR modulators

To investigate the binding properties of selected steroidal and non-steroidal ligands with human PR, we performed ensemble docking, by retaining the 5 highly repeating water molecule, with candidate crystal structures from each group. We selected the crystal structures with PDB IDs 2W8Y (from Group I), 2OVH (Group II), 3ZRB (Group III), and 3ZR7 (Group VI), since these proteins possessed best resolution among the groups (Table 1). The 2W8Y has a steroidal antagonist Mifepristone (RU486), but induced a destabilized agonistic conformation to PR upon binding. The 2OVH, 3ZRB and 3ZR7 consists of ligands such as asoprisnil, ORC and OR8 respectively.

A total of 51 steroidal and non-steroidal scaffold ligands with antagonist as well as SPRM properties were prepared for this docking studies (Table S1). When we effectuated rigid standard precision docking for all these 51 ligands with the 4 candidate receptor structures, it was noticed that ligands that are having steroidal scaffolds were not docked to 3ZRB and 3ZR7. The binding sites of these two proteins were accustomed to relatively small the non-steroidal ligands and induced fit conformational changes were observed in accordance with these non-steroidal ligands. We found that all the selected non-steroidal ligands were able to dock to all the 4 receptors.

MM-GBSA results showed that steroidal ligands having better binding energy than non-steroidal ligands as they have a greater number of hydrophobic interactions when compared to non-steroidal ligands. The binding energy for steroidal ligands ranges from -174.9 to 129.06 kcal/mol and non-steroidal from -125.39 to -43.93kcal/mol respectively (Table 2). Our structural analysis revealed that two different conformations (conf-1 and conf-2) are available for the LBD of PR, which is mainly due to the plasticity of the H12. The plasticity of H12 is largely determined by the ligands binds to the LBD. Steroidal ligands that are having bulky substitutions on the ‘C’ ring push the H12 away from the protein (conf-1). Conversely, steroidal ligands that are not having substitutions on ‘C’ ring doesn’t impart any conformational changes to H12(conf-2) (Figure 2). Similarly, the non-steroidal ligands that are relatively small in size were also not imparting conformational changes to H12. This conformational plasticity determines the recruitments of co-repressor and activators. Apart from the H12 plasticity, we could also see a small induced fit conformational changes to the binding site residues that are created by the respective ligands.

**Table 2:**
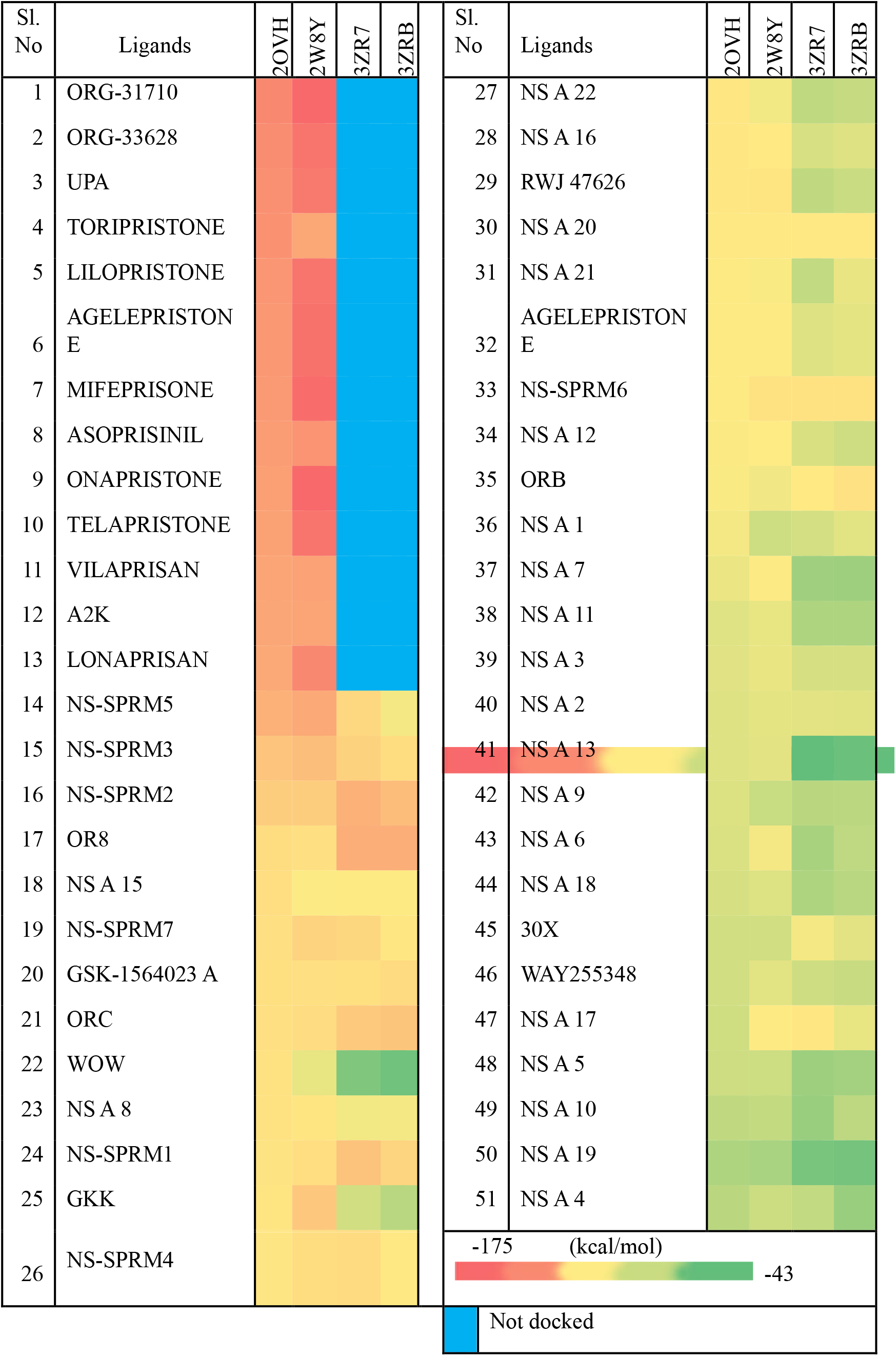
Binding energies of the 51 selected ligands (represented as heatmap) against four different receptors such as 2OVH, 2W8Y, 3ZR7 and 3ZRB. Colour codes in the respective heatmap is also shown.

Our molecular docking studies followed by binding energy calculations revealed that steroidal ligands that are having large substitutions on the ‘C’ ring always prefer the conf-1. We could see that the non-steroidal ligands were also easily accommodated to the conf-1. However, the non-steroidal ligands prefer Conf-2 as evident from the binding energy calculations. It was also noted that there is a decrease in the binding energies when ligands were docked in the absence of those repetitive water molecules, wat 4, emphasizing the importance of this highly repeating water molecule in ligand binding.

Though we have selected 51 ligands, we critically evaluated the binding properties of those ligands that are under clinical trial and has no complex crystal structures with PR. We selected 4 ligands, telapristone, vilaprisan, GSK-1564023A and WAY-255348 (Figure 1D-G). To explain its atomic level of interactions with PR, we executed ensemble docking followed by molecular dynamics simulations. As expected, the telapristone and vilaprisan were only docked to 2W8Y and 2OVH (Figure 4A and B). Similarly, the non-steroidal ligands GSK-1564023A and WAY-255348 were possessed better and similar binding energies towards 3ZR7 and 3ZRB (Figure 4C and D). Since our docking protocol was used rigid receptor, we performed the MD simulations to assess the binding stability. Our MD simulations showed that the proposed docking mode is highly stable and doesn’t deviate more than 1.0 Å (Figure 5).

**Figure 4:**
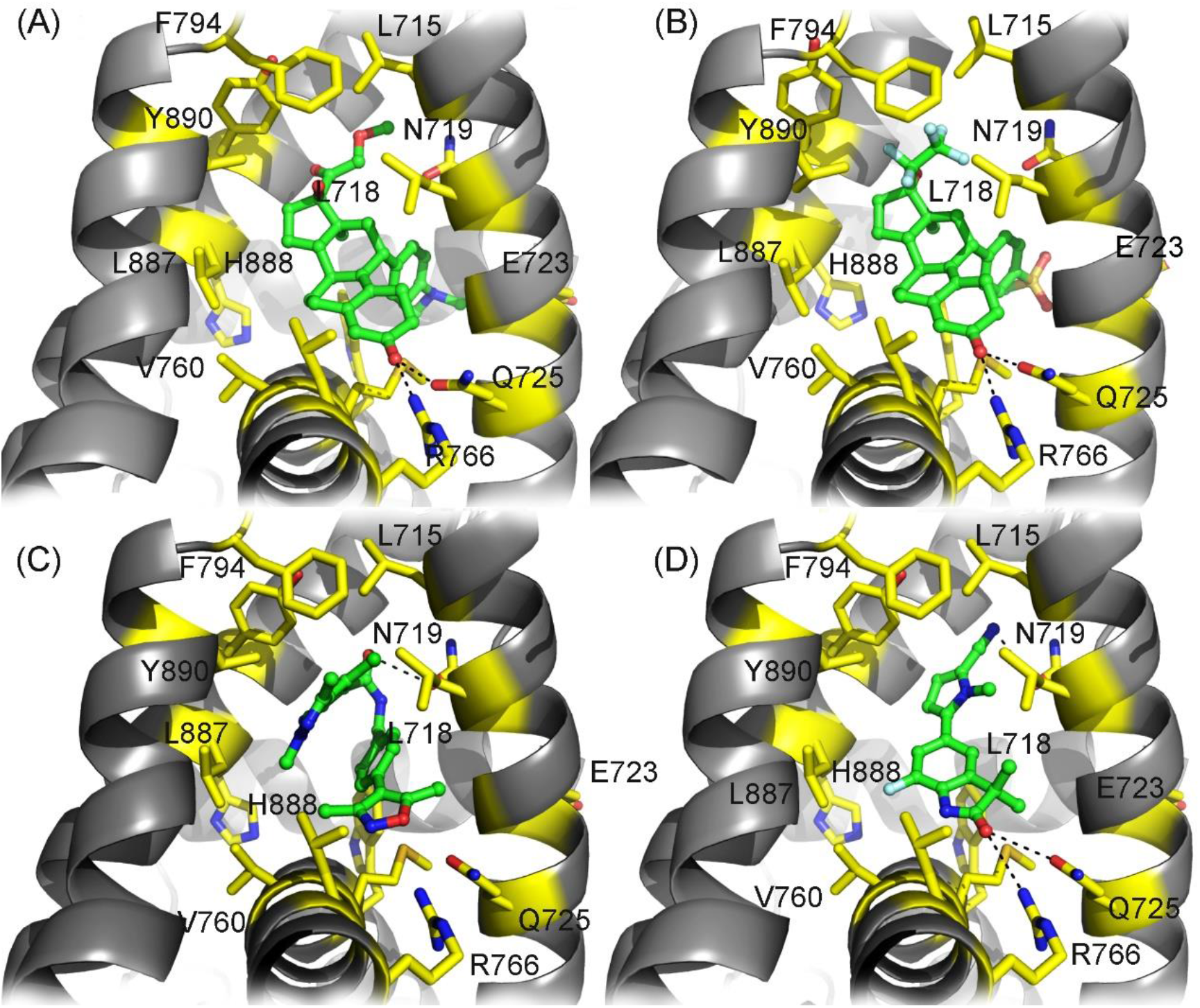
Binding modes of 4 selected PR modulators with the LBD of PR. Most energetically stable docked poses of selected steroidal ligands i.e., telapristone (A) and vilaprisan (B) in binding site of 2W8Y and 2OVH respectively. Similarly, the energetically stable binding modes of two selected non-steroidal ligands i.e., GSK-1564023A (C) and WAY-255348 (D) in the binding site of 3ZR7.

**Figure 5:**
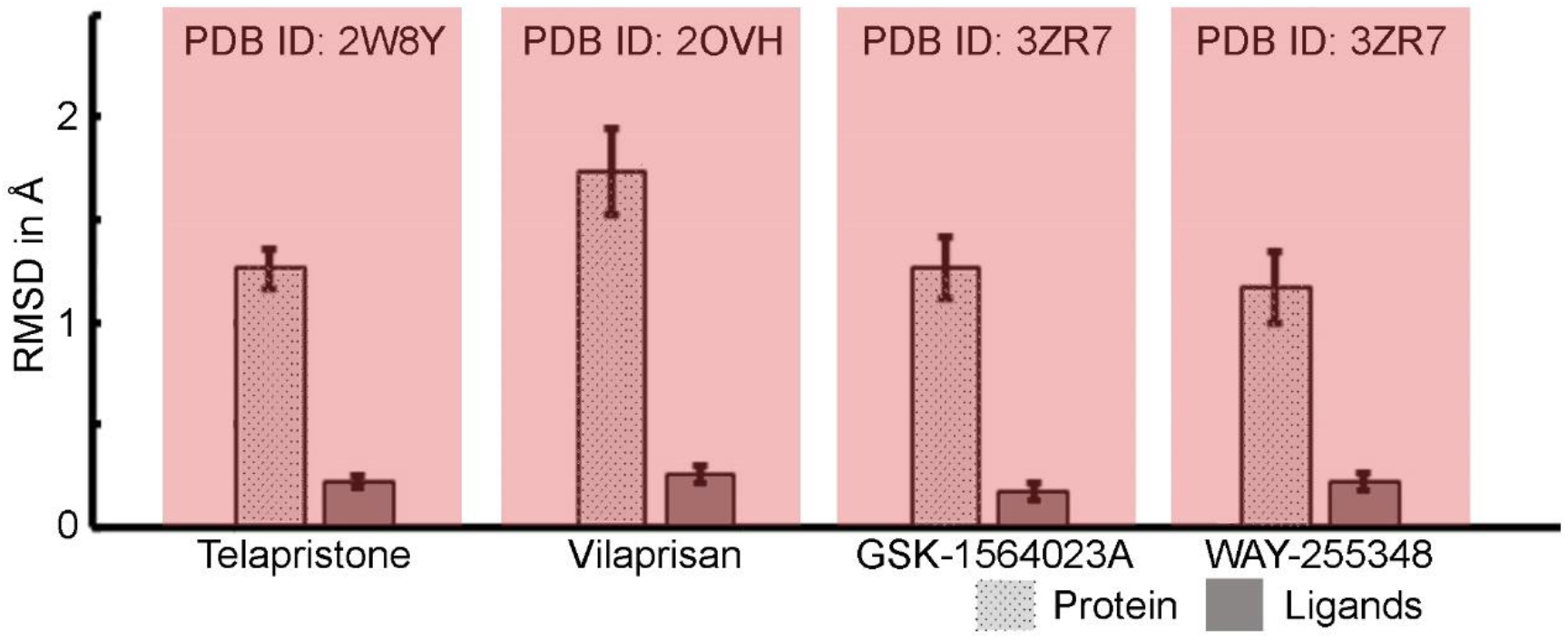
Root mean square deviation of protein and ligand complexes throughout 100 ns MD simulations. PDB ID of the selected PR structure and respective ligands were also mentioned.

### Telapristone adopted a similar binding mode as Mifepristone in 2W8Y

Upon docking of the Telapristone with 2W8Y (binding energy -163.22 kcal/mol), the steroid ring of the ligand exhibited a similar interaction pattern as that of Mifepristone [27,34,] (Figure 4A, 5 and S1A-C). The ketone group of the A-ring in Telapristone anchors to R766 and Q725 through hydrogen bonds. But, the 17β-hydroxyl on the D-ring does not forms a water-mediated hydrogen bond to N719, as in the case of RU486. The 17α-methoxyacetyl of Telapristone is accommodated in a cavity formed by L715, L718, F794, L797, M801, and Y890. Dimethylamine group of 11β substitution in Telapristone makes a strong cation-π interaction with W755 throughout the simulation time. This interaction allows more rigidity to the ligand and hence more stable in the closed conformation of PR. 11β substitution of Telapristone was 2.3Å apart from M909 of H12. Telapristone also forms hydrophobic contacts with F905 and V912.

### 11β pendant group of Vilaprisan plays an important role in its binding to 2OVH

Our docking studies also revealed that Vilaprisan binds strongly to 2OVH with a binding energy of -137.64 kcal/mol This strong binding is through a hydrogen bond from the ketone group of A ring to Q725 and R766 as similar to other steroidal ligands [30, 34] (Figure 4B). The 17β-hydroxyl on the D-ring forms a water-mediated hydrogen bond to N719, and this polar interaction is maintained more than 86% of stimulation time (Figure 5 and S2A-C). The oxygen atom of 11-methylsulfonylphenyl in the C ring of Vilaprisan makes a water bridged interaction with E723 which is persisted majority of stimulation time. Other interactions of Vilaprisan with 2OVH are water bridges with E723, Q725, C891, M759, R766 and hydrophobic interactions with L715, L718, L721, W755, L763, F778 and L797 respectively.

As in the case of Asoprisnil, the 11β pendant group of Vilaprisan is also occupied on the same space in the LBD near to the residue M909 [30]. To accommodate this pendant group within the ligand pocket, the loop region near H12 was pushed out to adopt a T-shaped binding pocket. i.e., Vilaprisan adopts conf-1 by pushing M909 to 16.5Å apart by the terminal methyl group of the bulky substituent at C ring of the ligand. This type of binding may render the SPRM activity of these ligands.

### GSK-1564023A shows a non-planar conformation in 3ZR7

In the docking studies of GSK-1564023A with 3ZR7, the binding modes of non-steroidal ligands exhibits similarities with that of steroidal ligands. Our docking studies pin pointed a direct H-bonding of oxygen atom in the isoxazole ring of GSK-1564023A to Q725 as seen in the case of Mifepristone where the ketone group is interacted with Q725. Nitrogen atom in the isoxazole adds stability through the conserved water molecule. We found that the following amino acids, i.e., R766, Q725 and M759 are involved in the water mediated interactions (Figure 4C, 5 and S3A-C). GSK-1564023A also directly bound to N719 through a H-bond via its amide nitrogen. This non-steroidal ligand is found to reside in a hydrophobic pocket lined with L718, L721, W755, M756, F778, F794, L797, L887, Y890, C891, F905 and M909. Additionally, a π-π stacking interactions of isoxazole ring with F778 and pyrazolopyridine with Y890 were observed.

The binding of GSK-1564023A is well tolerated within the same binding pocket as like in the steroidal ligands. Instead of planar conformation of steroid ligands, GSK-1564023A attained a C-shaped conformation in the LBD, as in the case of non-steroidal SPRM ‘OR8’ [29]. The Pyrazolopyridine ring of GSK-1564023A reaches the binding pocket as 17th position groups from steroids such as the 17α-methoxy acetyl group of mifepristone (27,34). Further analysis showed several van der waals interactions also strengthen the binding of ligand to PR.

### WAY-255348 mimic the binding mode of steroidal SPRM

The WAY-255348 showed almost planar conformation when compared to GSK-1564023A upon docked with 3ZR7. It has been found that the WAY-255348 is placed in a hydrophobic pocket lined by L718, M756, F778, Y890 and C891. Oxindole ring of WAY-255348 adopted almost similar orientation in the LBD as like A ring of Mifepristone. As seen in the steroidal ligands, the ketone group in this oxindole ring also makes water bridged interaction with R766, Q725 & M759 through the conserved water molecule. In the simulation studies, these interactions were found to maintain throughout the stimulation time. Additionally, the cyano group in the pyrrole ring of WAY-255348 interact with N719 through a hydrogen bond and this interaction persist throughout the stimulation time (Figure 4D, 5 and S4A-B).

## Conclusion

The SPRMs modulates PR action by allowing the coactivators and corepressors to interact with PR and thereby exhibiting mixed agonist and antagonist profiles. SPRMs are acting as an alternate therapeutic agent for the treatment of UF, endometriosis and breast cancer. In the case of UF treatment, the SPRMs display very similar effects on the reproductive system, blocking ovulation, suppressing bleeding and reducing the size of uterine leiomyoma. Though there are several SPRMs were put forwarded for the treatment of UF, due to the safety concern, they were withdrawn from the market. Especially the SPRMs such as UPA and ASO were exhibited unfavorable endometrial changes in UF patients and liver toxicity respectively. In the present scenario there is a dire need for new SPRMs with high efficacy and little side effects. To contribute towards the identification of novel SPRMs, we put forwarded the binding modes of two steroidal (telapristone and vilaprisan) and non-steroidal ligands (GSK-1564023A and WAY-255348) with the ligand binding domain of PR through molecular modeling. To the best of our knowledge the binding modes of these ligands were not solved either through experimental or by computational ways. Our extensive structural analysis identified five highly repeating water molecules. One of these water molecules is contributing towards the ligand binding and stability as it is having a direct interaction with the ligands. Our ensemble docking suggested that steroidal ligands bind more strongly to human PRs than non-steroidal ligands. Our docking studies demonstrated that the steroidal ligands that are having bulky substitutions on the ‘c’ ring can only recognize open conformation of the helix-12, whereas the non-steroidal ligands can bind to both open and close conformations of helix-12. Binding energies of non-steroidal ligands towards the closed conformation is high when compared to that of open conformation as they make more interactions in the closed conformations. Hence, a high throughput virtual screening campaign that leads to the identification of SPRMs could be enriched by including the information about the invariant water molecules (especially Wat-5). Similarly, selecting an appropriate receptor structure (either open or close conformations of helix-12) could also enrich the studies. We anticipate that our studies would certainly be useful for the identification of novel lead PR modulators with improved efficacy and safety profiles.

## Acknowledgement

The authors are thankful for the computational facilities at Laboratory for Structural Biology, RIKEN, Japan. The financial assistance from Department of Health Research, Govt. of India (Project number R.12015/01/2021-HR/E-Office:8116780) is gratefully acknowledged.

