## Supplemental tables and figures for "Structural analysis and ensemble docking revealed the binding modes of selected progesterone receptor modulators"

**Figure S1:** Conformational changes of 2W8Y-telapristone complex throughout 100 ns MD simulations. (A) RMSF of the protein. (B) torsion angle profiles of telapristone and (C) number of protein ligand contacts (≤ 4Å) throughout simulations.

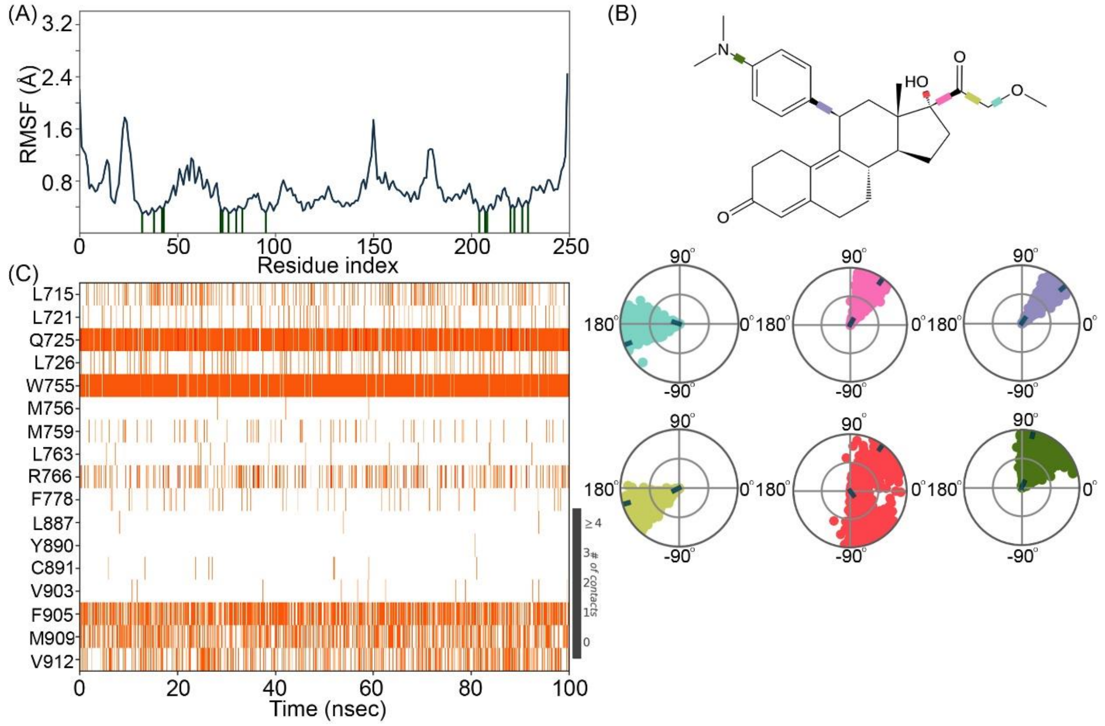


**Figure S2:** Conformational changes of 2OVH-vilaprisan complex throughout 100 ns MD simulations. (A) RMSF of the protein. (B) torsion angle profiles of telapristone. (C) number of protein ligand contacts (≤ 4Å) throughout simulations.

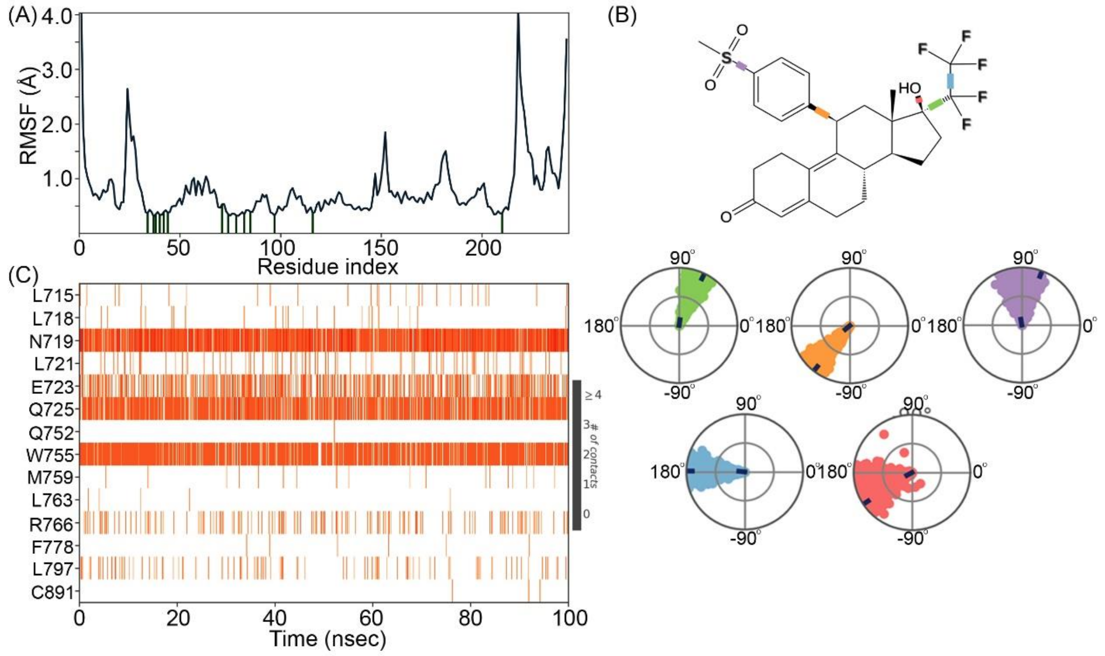


**Figure S3:** Conformational changes of 3ZR7-GSK-1564023A complex throughout 100

ns MD simulations. (A) RMSF of the protein. (B) torsion angle profiles of GSK-1564023A. (C) number of protein ligand contacts (≤ 4Å) throughout simulations.

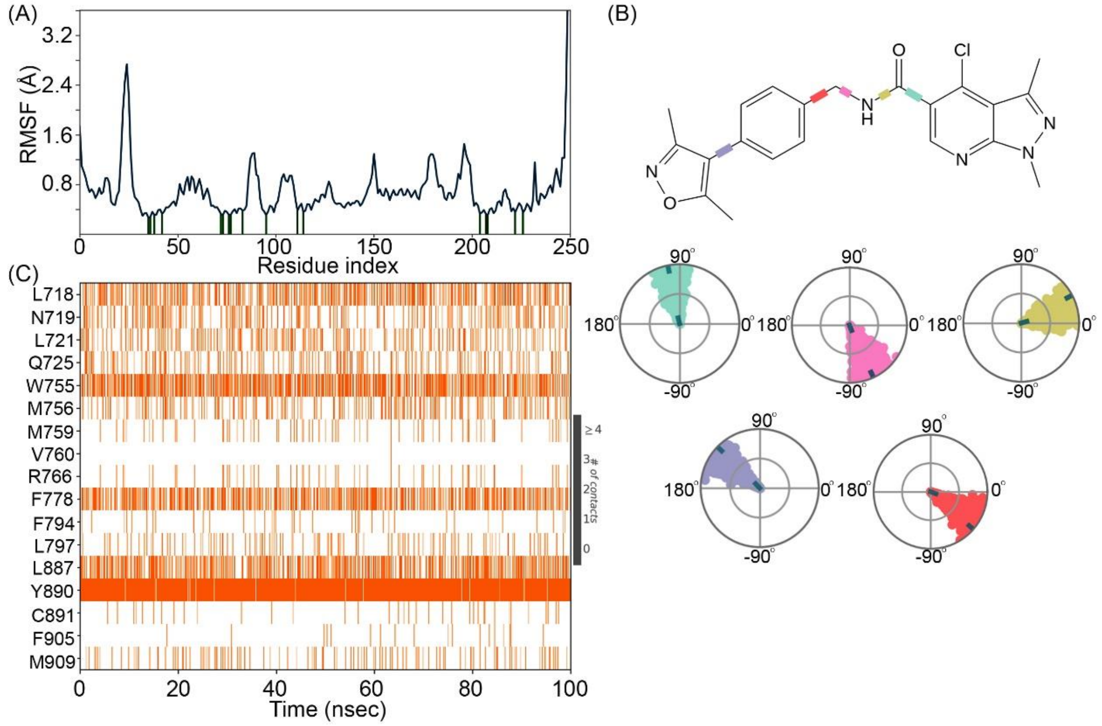


**Figure S4:** Conformational changes of 3ZRB-WAY-255348 complex throughout 100 ns MD simulations. (A) RMSF of the protein. (B) torsion angle profiles of WAY-255348 and (C) number of protein ligand contacts (≤ 4Å) throughout simulations.

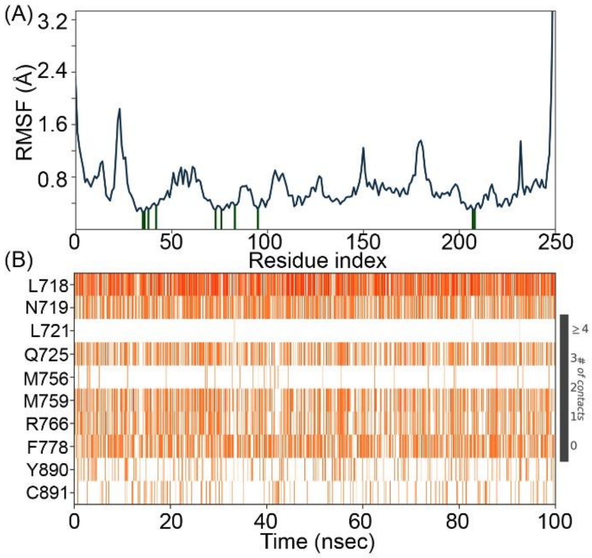

**Table S1:** Names and smiles of the 51 known PR antagonists used for the docking studies. Out of theses 51 ligands, 13 has steroidal and 38 were in non-steroidal scaffold. NS-SPRM and NSA indicates Non-steroidal SPRMs and Non-steroidal Antagonists respectively.

| **Human PR Ligands** | | | **SMILE FORMAT** |
|---|---|---|---|
| Antagonist | Steroids | Mifepristone | CC#C[C@]1(O)CC[C@H]([C@@]12C)[C@H]3C(=C4C(CC3)=CC(=O)CC4)[C@H](C2)c5ccc(N(C)C)cc5 |
| | | ORG-31710 | C1CCO[C@@]12[C@]3(C)[C@@H](CC2)[C@H]4C(=C5C([C@@H](C4)C)=CC(=O)CC5)[C@H](C3)c6ccc(N(C)C)cc6 |
| | | ORG33628 | C=C1CCO[C@@]12[C@]3(C)[C@@H](CC2)[C@H]4C(=C5C(CC4)=CC(=O)CC5)[C@H](C3)c6ccc(cc6)C(=O)C |
| | | AGLEPRISTONE | C/C=C\[C@]1(O)CC[C@H]([C@@]12C)[C@H]3C(=C4C(CC3)=CC(=O)CC4)[C@H](C2)c5ccc(N(C)C)cc5 |
| | | LONAPRISAN | FC(F)(F)C(F)(F)[C@]1(O)CC[C@H]([C@@]12C)[C@H]3C(=C4C(CC3)=CC(=O)CC4)[C@H](C2)c5ccc(cc5)C(=O)C |
| | | LILOPRISTONE | OC/C=C\[C@]1(O)CC[C@H]([C@@]12C)[C@H]3C(=C4C(CC3)=CC(=O)CC4)[C@H](C2)c5ccc(N(C)C)cc5 |
| | | TORIPRISTONE | CC#C[C@]1(O)CC[C@H]([C@@]12C)[C@H]3C(=C4C(CC3)=CC(=O)CC4)[C@H](C2)c5ccc(cc5)N(C)C(C)C |
| | | ONAPRISTONE | OCCC[C@@]1(O)CC[C@H]([C@]12C)[C@H]3C(=C4C(CC3)=CC(=O)CC4)[C@H](C2)c5ccc(N(C)C)cc5 |
| | Non-Steroids | RWJ-47626 | FC(F)(F)c(c1)c(Cl)ccc1C2=NN(CCC2)S(=O)(=O)c3c(Cl)ccc(c3)C(F)(F)F |
| | | WAY-255348 | CC1(C)C(=O)Nc(c12)c(F)cc(c2)-c(n3C)ccc3C#N |
| | | GSK-1564023A | Cc1nn(C)c(c12)ncc(c2Cl)C(=O)NCc3ccc(cc3)-c4c(C)onc4C |
| | | ORB | Clc1ncc(F)cc1-c(cc2)ccc2[C@@H](C)NS(=O)(=O)c3c(C)onc3C |
| | | ORC | Cc1noc(C)c1S(=O)(=O)N[C@H](C)c2ccc(cc2)-c3c(C)onc3C |
| | | NSA1 | ON(O)c(c1)cccc1- |

| | | | |
|---|---|---|---|
| | | | c(cc2)cc(c23)n(c(=O)[nH]3)C(C)C |
| | | NSA2 | ON(O)c(c1)cccc1-c(cc2)cc(c23)sc(=O)[nH]3 |
| | | NSA3 | c1sc(C#N)cc1-c(cc2)cc(c23)n(C)c(=O)[nH]3 |
| | | NSA4 | [nH]1c(=O)sc(c12)cc(cc2)-c3ccc(C#N)cc3 |
| | | NSA5 | N#Cc(c1)cc(F)cc1-c(cc2)cc(c23)C(C)(C)C(=O)N3 |
| | | NSA6 | N#Cc(c1)cc(F)cc1-c(c2)ccc(NC3=O)c2C34CCCC4 |
| | | NSA7 | N#Cc(c1)cc(F)cc1-c(c2)ccc(NC3=O)c2C34CCCCC4 |
| | | NSA8 | c1c(Cl)cccc1-c(c2)ccc(N[S@@]3=O)c2C34CCCCC4.C=O |
| | | NSA9 | N#Cc(c1)c(F)ccc1-c(cc2)cc(c23)C(C)(C)OC(=O)N3 |
| | | NSA10 | c1oc(C#N)cc1-c(cc2)cc(c23)C(C)(C)OC(=O)N3 |
| | | NSA11 | c1c(F)cc(C#N)cc1-c(cc2)cc(c23)C(C)(C)OC(=O)N3 |
| | | NSA12 | c1sc(C#N)cc1-c(cc2)cc(c23)C(C)(C)OC(=O)N3 |
| | | NSA13 | N#Cc(c1)cc(F)c(c12)OCc3c2ccc4c3[C](C)(C)=CC(=O)C4 |
| | | NSA14 | c1ccc(Cl)cc1-c(cn2)cc(c23)C(C)(C)OC(=O)N3 |
| | | NSA15 | c1c(Cl)cccc1-c(cn2)cc(c23)[C@](C)(N(C)C(=O)N3)C4CC4 |
| | | NSA16 | c1c(Cl)cccc1-c(cc2)cc(c23)C(C)(C)CC(=O)N3 |
| | | NSA17 | CC(C)(C)CN(CC)c(cc1)cc(c12)c(C(F)(F)F)cc(=O)[nH]2 |
| | | NSA18 | C1CCCCC1c(cc2)cc(c23)c(C(F)(F)F)cc(=S)[nH]3 |
| | | NSA19 | c1c(F)cc(C#N)cc1-c(cc2)cc(c23)c(C(F)(F)F)cc(=S)[nH]3 |
| | | NSA20 | C1CCCC=C1c(cc2)cc(c23)C(C)(C)OC(=O)N3 |
| | | NSA21 | C1CCCC=C1c(cc2)cc(c23)C(C)(C)OC(=S)N3 |

| | | | |
|---|---|---|---|
| | | NSA22 | C1CCC(=O)C=C1c(cc2)cc(c23)C(C)(C)OC(=S)N3 |
| Selective PR Modulators | Steroids | ASOPRISNIL | COC[C@]1(OC)CC[C@H]([C@@]12C)[C@H]3C(=C4C(CC3)=CC(=O)CC4)[C@H](C2)c5ccc(cc5)\C=N\O |
| | | A2K | C1CC1C(=O)[C@H]2[C@H](C=C)C[C@H]([C@@]23C)[C@H]4C(=C5C(CC4)=CC(=O)CC5)[C@H](C3)c6ccc(cc6)-c7cccnc7 |
| | | TELAPRISTONE | COCC(=O)[C@@]1(O)CC[C@H]([C@@]12C)[C@H]3C(=C4C(CC3)=CC(=O)CC4)[C@H](C2)c5ccc(N(C)C)cc5 |
| | | Ulipristal Acetate | O=C(C)O[C@]1(C(=O)C)CC[C@H]([C@@]12C)[C@H]3C(=C4C(CC3)=CC(=O)CC4)[C@H](C2)c5ccc(N(C)C)cc5 |
| | | VILAPRISAN | O=S(=O)(C)c(cc1)ccc1[C@@H](C2)C(=C3C(CC4)=CC(=O)CC3)[C@@H]4[C@@H]([C@]25C)CC[C@@]5(O)C(F)(F)C(F)(F)F |
| | Non-Steroids | 30X | CN(C)C(=O)[C@H](C)N(CC(F)(F)F)c1cc(C(F)(F)F)c(C#N)cc1 |
| | | OR8 | Cn1nc(C)c(c1Cl)S(=O)(=O)NCc2ccc(cc2)-c3c(C)onc3C |
| | | GKK | N#Cc(cc1)c(Cl)cc1N([C@@H](CC2)CN2C)Cc3c(C(F)(F)F)cccc3 |
| | | WOW | N#Cc(cc1)c(Cl)cc1N([C@@H](CC2)CN2S(=O)(=O)C)Cc3c(C)cccc3 |
| | | NS-SPRM1 | ON(O)c1cc(C)c(cc1)/N=C(\CC2CC2)N([C@H](C)CC)C3CCCC3 |
| | | NS-SPRM2 | ON(O)c1cc(C)c(cc1)/N=C2/N([C@H](C)CC)[C@H](CS2)CC(C)C |
| | | NS-SPRM3 | ON(O)c1cc(C)c(cc1)/N=C2/N(C(=O)[C@H](S2)C(C)C)CC(CC)CC |
| | | NS-SPRM4 | ON(O)c(c1)ccc(c1C)/N=C(OC2)/N(C23CCCC3)C4CCCCC4 |
| | | NS-SPRM5 | ON(O)c(c1)ccc(c1C)N(C(CCC)CCC)C2=N[C@H](CS2)CC(C)C |
| | | NS-SPRM6 | c1cc(N(O)O)ccc1-c(cc2)cc(c23)C(C)(C)[C@H](N3)C |
| | | NS-SPRM7 | c1cc(N(O)O)ccc1-c(c2)ccc(N[C@@H]3C)c2C34CCCCC4 |
